# Chromone scaffold–mediated reprogramming of the epithelial–mesenchymal transition prevents fibrosis

**DOI:** 10.1101/106591

**Authors:** Han-Soo Kim, Jun-Hwan Kim, Young-Min Yoon, Moon Kee Meang, Ji Yong Lee, Tae Hee Lee, Ik-Hwan Kim, Byung-Soo Youn

**Author notes:** These authors contributed equally to this work (Han-Soo Kim, Jun-Hwan Kim). To whom correspondence should be addressed: OsteoNeuroGen, Inc. Ace High-end Tower 9th 233, Gasandigital-1-ro, Geumcheon-gu, Seoul, 08501, Republic of Korea.

## Abstract

Fibrotic diseases are major causes of morbidity and mortality, and the epithelial–mesenchymal transition (EMT) plays a central role in the development of tissue/organ fibrosis. We discovered that eupatilin, a member of the chromone scaffold (CS)-containing compounds found ubiquitously in the plant kingdom, completely reversed fibrogenesis *in vitro* and substantially ameliorated bleomycin-induced lung fibrosis (BILF). Furthermore, eupatilin-induced growth arrest and morphological changes in primary fibroblasts derived from a patient with idiopathic pulmonary fibrosis (IPF). To better understand fibrosis, we established a mouse hepatic stellate cell (HSC) line that was robustly differentiated into myofibroblasts upon treatment with TGFβ. HSC-derived fibrogenesis was completely blocked by eupatilin, which caused dramatic morphological changes while inhibiting expression of EMT-related genes. The chemical groups linked to the 2^nd^ carbon (C2), C3, C6, and C7 on the CS of eupatilin were essential for its anti-fibrogenic effects. Unlike eupatilin, pirfenidone failed to block HSC fibrogenesis and did not affect the morphology of HSCs or lung fibroblasts. Although pirfenidone affected local production of TGFβ, as reflected by a reduction in the TGFβ level in lung lysates of BILF model mice, eupatilin is likely to act via a different therapeutic mechanism. In particular, eupatilin had greater anti-fibrotic capacity and EMT-inhibitory activity and significantly attenuated the phosphorylation of Erk by TGFβ. Based on the interactome, *Integrinβ3* seems to be a major player in integration of TGFβ signaling into the eupatilin-mediated anti-fibrosis. Our findings suggest that combinatorial use of eupatilin and pirfenidone may augment the therapeutic efficacy of IPF treatment.

## Introduction

Fibrosis is a complex disease state in which elevated proliferation of myofibroblasts is accompanied by overproduction of extracellular matrix (ECM) components (Bonnans et al., 2014). Two fibrotic conditions, idiopathic pulmonary fibrosis (IPF) and liver fibrosis preceded by non-alcoholic steatohepatitis (NASH), affect many people around the world, and thus represent an urgent medical need. To date, however, no curative drugs for these conditions have become available (Fallowfield, 2015; Sgalla et al., 2016). Two small-molecule anti-fibrotic drugs, pirfenidone (Esbriet) and nintedanib (Ofev), have recently been approved for IPF (King and Nathan, 2015), but their efficacies and side effects must be improved to successfully treat this disease. Moreover, while nintedanib is an angio-kinase inhibitor, mode of action of pirfenidone exerting their anti-fibrotic activities remain to be elucidated.

Flavones are members of the polyphenol family, a group of >10,000 compounds predominantly found in the plant kingdom (Dixon and Pasinetti, 2010). In general, these phytochemicals protect plants from radiation damage (Stapleton and Walbot, 1994). Due to their anti-oxidant and anti-inflammatory potentials, flavones have long been used to treat inflammatory diseases such as arthritis and asthma (Leyva-López et al., 2016). Chromone, 1,4-benzopyrone-4-one, is a central chemical scaffold constituting flavones (Emami and Ghanbarimasir, 2015), and the chromone derivatives (CDs) are a diverse family based on branching chemical residues coupled to this core chemical framework (Gacche et al., 2015). We recently reported that eupatilin, a CD from an *Artemisia* species, dramatically inhibits osteoclastogenesis (Kim et al., 2015) and downregulates multiple genes associated with the epithelial–mesenchymal transition (EMT). Specifically, 24 of the 50 top genes differentially regulated by eupatilin are associated with the EMT, prompting us to hypothesize that eupatilin can prevent fibrosis (Figure S1).

Here, we show that eupatilin can block induction of EMT or cell growth arrest and alter cell morphology, thereby substantially ameliorating lung fibrosis. This observation opens a door as a powerful new therapeutic modality for treating IPF.

## Results and discussion

### Eupatilin has potent anti-fibrogenic capacity

We first investigated whether TGFβ-mediated fibrogenesis could be inhibited by CDs, particularly eupatilin, in human lung fibroblasts derived from IPF patients (hereinafter DHLFs: Diseased Human Lung Fibroblasts) (Huang et al., 2014). Compared with TGFβ alone, costimulation of DHLFs with TGFβ plus eupatilin resulted in significant growth arrest and morphological change (Figure 1A and S2A). Currently, pirfenidone and nintedanib are the only targeted drugs approved for use in treating IPF (du Bois, 2010). Two different concentrations of nintedanib completely blocked myofibroblastogenesis, whereas pirfenidone had no effect (Figure S2B).

**Figure 1.**
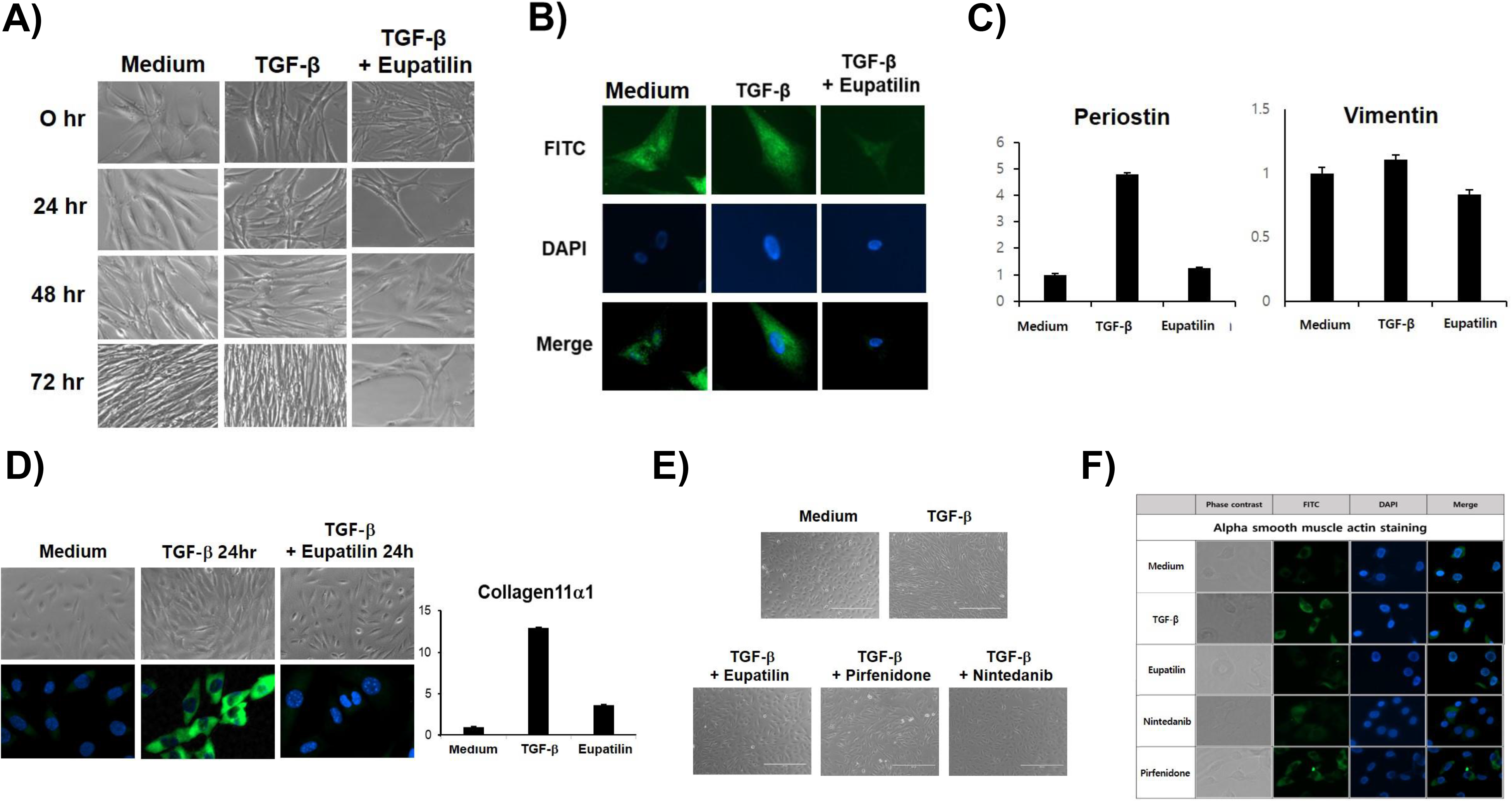
Inhibition of TGFβ-induced fibrogenesis by eupatilin. (A) DHLFs (5 × 10^6^ cells) were seeded in fibroblast growth medium (FBM), grown overnight, and then stimulated with control medium, TGFβ (5 ng/ml), or TGFβ plus eupatilin (50 μM) for 72 hr. Morphological change was monitored by phase-contrast microscopy at 40× magnification. (B) DHLFs were stimulated as described above, and ICC was performed using antibody against mouse α-smooth muscle actin. Nuclei were detected by DAPI staining. (C) Total RNA was isolated from DHLFs stimulated with control medium, TGFβ, or TGFβ plus eupatilin for 48 hr, and then subjected to real-time PCR analysis using primers against *periostin* or *vimentin*. Standard deviations were calculated from the results of three independent PCRs. (D) ONGHEPA1 cells were stimulated with TGFβ (5 ng/ml) or TGFβ plus eupatilin (50 μM) for 24 hr, and morphological changes were monitored under a light microscope. ICC was performed using anti-mouse α-smooth muscle actin. (E) Total RNAs were isolated from ONGHEPA1 cells stimulated with medium, TGFβ, or TGFβ plus eupatilin for 48 hr, and then subjected to real-time PCR analysis using primers against *collagen11α1*. (F) ONGHEPA1 cells were grown and stimulated with medium, eupatilin, pirfenidone, or nintedanib in the presence or absence of TGFβ for 24 hr, and then anti-fibrogenesis was monitored. (G) ONGHEPA1 cells were stimulated as described above, and ICC was performed using antibody against mouse α-smooth muscle actin. Nuclei were detected by DAPI staining.

To examine the influence of these compounds on the EMT, we performed immunocytochemistry (ICC) to detect expression of α-smooth muscle actin (αSMA), a marker of EMT induction. DHLFs, presumably preconditioned in IPF patients, had a myofibroblast-like morphology *in vitro* and could intrinsically express αSMA, which was slightly upregulated by TGFβ (Figure 1B). Co-stimulation of DHLFs with eupatilin and TGFβ, however, virtually eliminated αSMA expression (Figure 1B). Among the EMT-associated genes upregulated in DHLFs are those encoding vimentin and periostin; the latter is an important regulator of ECM biogenesis that also serves as a ligand for integrin αVβ3 (Morra and Moch, 2011). TGFβ substantially increased expression of both genes, but eupatilin negated their upregulation (Figure 1C).

A549 cells are a human lung adenocarcinoma cell line of alveolar basal epithelial cell origin that have been used for analysis of tumor metastasis and induction of EMT (Kasai et al., 2005). TGFβ rendered A549 cells somewhat blastogenic and caused them to significantly upregulate *Snail* and *Vimentin*, but these effects were substantially attenuated by eupatilin (Figure S3). Moreover, cell morphology was changed dramatically in that the resultant cells became rounder, suggesting that eupatilin could reorganize cytoskeletal structures. Taken together, these data suggest that eupatilin inhibits proliferation and drives morphological change of DHLFs, thereby negatively regulating EMT. In previous work, we demonstrated that eupatilin does not promote cell death (Kim et al., 2015).

Myofibroblasts play a pivotal role in collagen deposition in fibrotic tissues. Differentiation of several progenitor cell types has been observed in lung tissues, but reports of *in vitro* lung fibrogenesis remain limited (Pretheeban et al., 2012). To develop an efficient cell-based fibrogenesis assay, we established a mouse hepatic stellate cell (HSC) line, ONGHEPA1 (Figure S4). HSCs are widely appreciated for their unique roles in initiation and perpetuation of liver fibrosis (Gabele et al., 2003). ONGHEPA1 had a mesenchymal morphology, expressed GATA4 but not cytokeratin 18 (Figure S4A and S4B), and expressed CD29, CD44, and CD106 on the cell surface (Figure S4C), strongly indicating that they are of mesenchymal origin. Based on these features, we concluded that ONGHEPA1 is a mouse primary HSC line.

When ONGHEPA1 cells were stimulated with TGFβ, full differentiation into myofibroblasts occurred within 12–24 hr, but eupatilin completely inhibited TGFβ-induced fibrogenesis at concentrations of 50 μM or higher (Figure S5A). However, pretreatment of ONGHEPA1 with eupatilin for 1–6 hr did not diminish fibrogenesis, suggesting that TGFβ-elicited signal transduction in combination with eupatilin is necessary to blunt fibrogenesis (Figure S5B). Eupatilin did not alter phosphorylation of Smad2 or Akt, but it significantly inhibited phosphorylation of Erk and marginally inhibited Smad3 after 12 hr of treatment; Erk phosphorylation recovered slightly at 24 hr (Figure S6), an observation we do not currently understand. Erk (Xie et al., 2004) or Smad3 (Willis and Borok, 2007) has been reported to play a pivotal role in TGFβ-mediated induction of the EMT.

PDGFR signaling has also been implicated in fibrogenesis (Wu et al., 2013). However, when ONGHEPA1 cells were subjected to differentiation, PDGF was incapable of driving differentiation in the presence of eupatilin (Figure S7A and S7B). ICC experiments clearly demonstrated that strong αSMA immunoreactivity induced by TGFβ stimulation was eliminated by eupatilin (Figure 1D) and the same pattern held true for real-time expression of *Col11α1* (Figure 1D).

Nintedanib is an angio-kinase inhibitor that antagonizes TGFβR, FGFR, and PDGFR, but the mode of action (MOA) of pirfenidone remains elusive (Wollin et al., 2015). When we stimulated cells with TGFβ in the presence of one of these drugs, or eupatilin, we observed that eupatilin and nintedanib effectively curbed fibrogenesis by ONGHEPA1 cells, as in the case of DHLFs, albeit with slight morphological differences between the two drugs (Figure 1E). On the other hand, pirfenidone could not inhibit differentiation, implying that the MOAs of eupatilin and pirfenidone may differ through this *in vitro* differentiation setup. The degree of inhibition of αSMA mirrored the anti-fibrogenic effects (Figure 1F). Taken together, these observations demonstrate that a chromone scaffold (CS)-containing compound, eupatilin, is a potent anti-fibrotic drug that can inhibit fibrogenesis during differentiation of HSCs into myofibroblasts largely by reprogramming EMT, leading to cell shape change. This conclusion in conjunction with the eupatilin-associated growth arrest and morphological changes in DHLFs prompted us to investigate whether eupatilin could exert a therapeutic effect on IPF. For this purpose, we used the mouse bleomycin-induced lung fibrosis (BILF) model.

### Eupatilin significantly improved BILF

Eupatilin or pirfenidone at 100 mg/kg (mpk) was orally administered twice per day, followed by one intratracheal administration of bleomycin; daily oral drug intake continued for the next 14 days (Figure 2). At 100mpk, the molar concentrations of pirfenidone is 2 fold higher than eupatilin. Oral administration of pirfenidone over 100mpk exhibited hepatotoxicity such that we simply selected 100mpk for comparing therapeutic capacity between these two drugs. Each drug administration group consisted of eight mice. We observed significant reductions in the levels of soluble collagen and hydroxyproline as well as changes in collagen histomorphometry in the drug-treated animals (Figure 2A, 2B, and 2C). Interestingly, the largest reduction in TGFβ production occurred in the pirfenidone-treated mice (Figure 2D), suggesting that the MOA of pirfenidone relates to downregulation of TGFβ production, presumably from immune cells. Fibrotic tissues are characterized by notable collagen deposition, along with proliferation or infiltration of myofibroblasts (Wynn, 2008). To examine these processes, we performed H&E staining in such a way as to visibly stain proliferative and infiltrated cells and used Sirius red to stain regions of collagen deposition. As seen in Figure 2E, bleomycin-treated lungs harbored the largest number of H&E- and Sirius red–stained cells, and pirfenidone treatment weakened the intensities of both stains. More intriguingly, eupatilin-treated lungs contained the smallest number of fibrotic cells and very pale collagen staining, suggesting that eupatilin may seem to have more potent cell regeneration potential than pirfenidone (Figure 2E).

**Figure 2.**
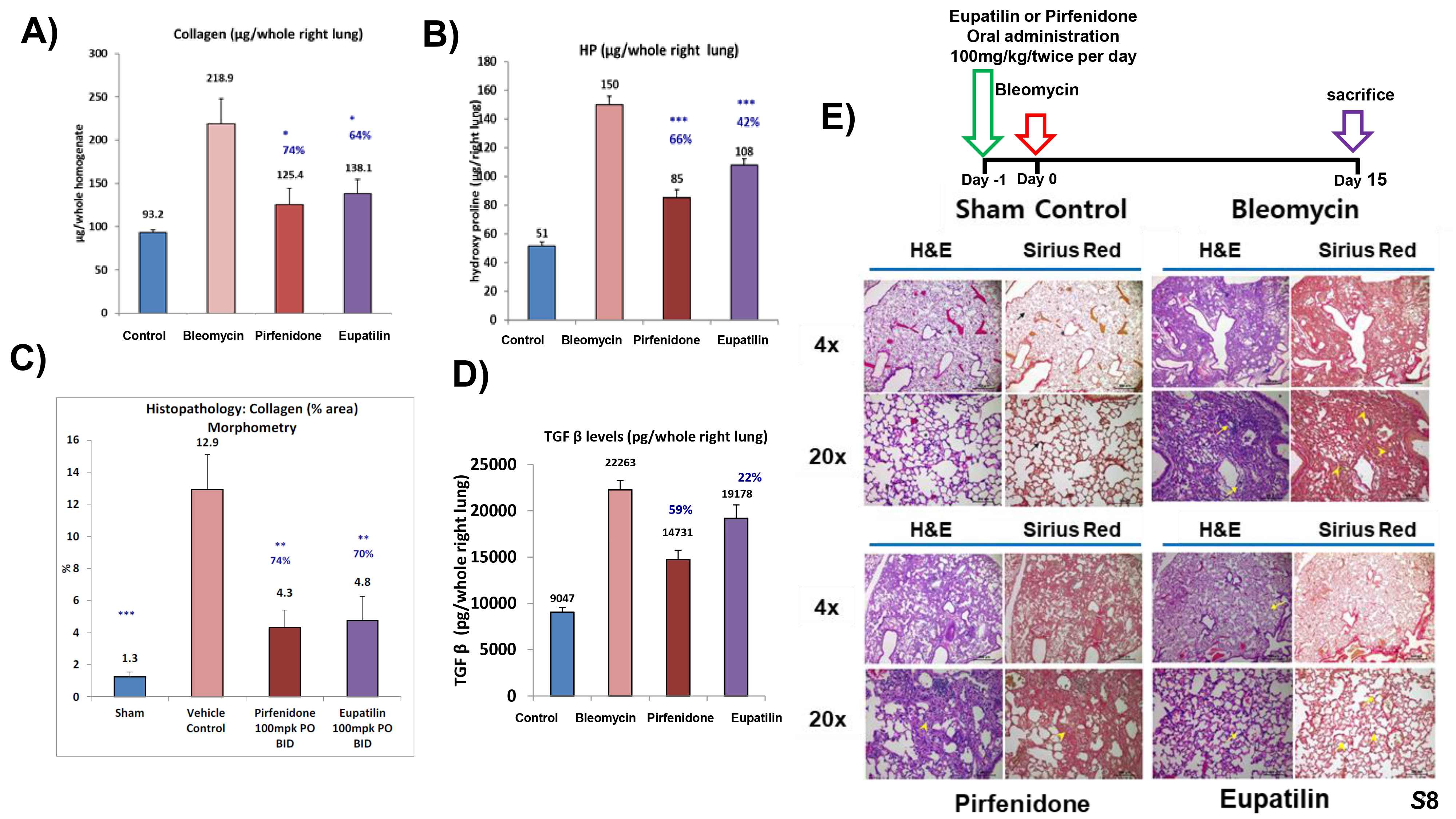
Anti-fibrotic efficacy of eupatilin. One intratracheal injection of bleomycin gave rise to fulminant fibrosis of the lungs in all mice. Vehicle, pirfenidone (100mpk), or eupatilin (100mpk) was orally administered twice per day for 14 days, and then lung tissue lysates and plasma were prepared. (A) Soluble collagen from whole right lung was measured. * denotes statistical significance (p<0.05, Student’s *t*-test). (B) Levels of hydroxyproline were measured and subjected to *t*-test. **, p<0.05 via *t*-test. (C) Collagen histomorphometry was examined and compared. Statistical significance was calculated by Student’s *t*-test. **, p<0.01; ***, p<0.001. (D) TGFβ in whole right lung was measured by ELISA. (E) Paraffin sections of lung tissues were stained with H&E or Sirius Red and examined using phase-contrast microscopy at 4× or 20× magnification. Infiltrated cells and collagen deposition were detected.

Next, to substantiate the anti-fibrotic potential of eupatilin, we examined the dose dependence of its effects. At doses of 100 and 200mpk, but not a lower dose of 50mpk, we observed a statistically significant improvement in the levels of soluble collagen and hydroxyproline. Overall, pirfenidone 100mpk had less of an inhibitory effect on collagen histomorphometry, soluble collagen, and hydroxyproline than eupatilin 200mpk, and this difference was also statistically significant (Figure 3A and 3B). Notably in this regard, the molar concentrations of eupatilin 200mpk is similar to those of pirfenidone 100mpk, suggesting that eupatilin is a more potent anti-fibrotic agent. As before, pirfenidone 100mpk was associated with a lesser amount of TGFβ production in comparison with eupatilin 200mpk (Figure 3C), again suggesting that the MOA of pirfenidone involves decreasing the production of TGFβ in the lung. Nevertheless, eupatilin 100mpk or 200 mpk was also able to significantly inhibit TGFβ production. Eupatilin has been reported to be efficiently absorbed into the upper GI tract and appears in plasma within 1 hr of ingestion, and is also readily detected in lung tissue lysates (Ji et al., 2004). We found that glucuronide-eupatilin was the major metabolite in plasma and excreted in urines (Figure S8A) whereas free eupatilin remained predominated in lung tissues. No glucouronide-eupatilin was detected by NMR. This observation suggests that orally administered eupatilin is rapidly disseminated to the lung via glucuronidation presumably occurring in the liver followed by deconjugation of the glucuronide moiety in the lung. Lung tissues express high levels of deconjugating enzymes (Strauss and Barbieri, 2014). Eupatilin existed in the lung exclusively as the free form but at concentrations at least 50-fold lower than those of pirfenidone in terms of molar concentrations, suggesting that eupatilin is an extremely potent drug against lung fibrosis as seen in Figure S8B. Pirfenidone is hepatotoxic at high concentrations (Verma et al., 2017), whereas the toxicity of eupatilin remains unknown. To explore this issue, we performed a full set of liver function tests using mouse plasma. As shown in Figure S9, we detected no hepatotoxicity from mice treated with either pirfenidone 100mpk or eupatilin200mpk, suggesting that eupatilin does not elicit hepatotoxicity.

**Figure 3.**
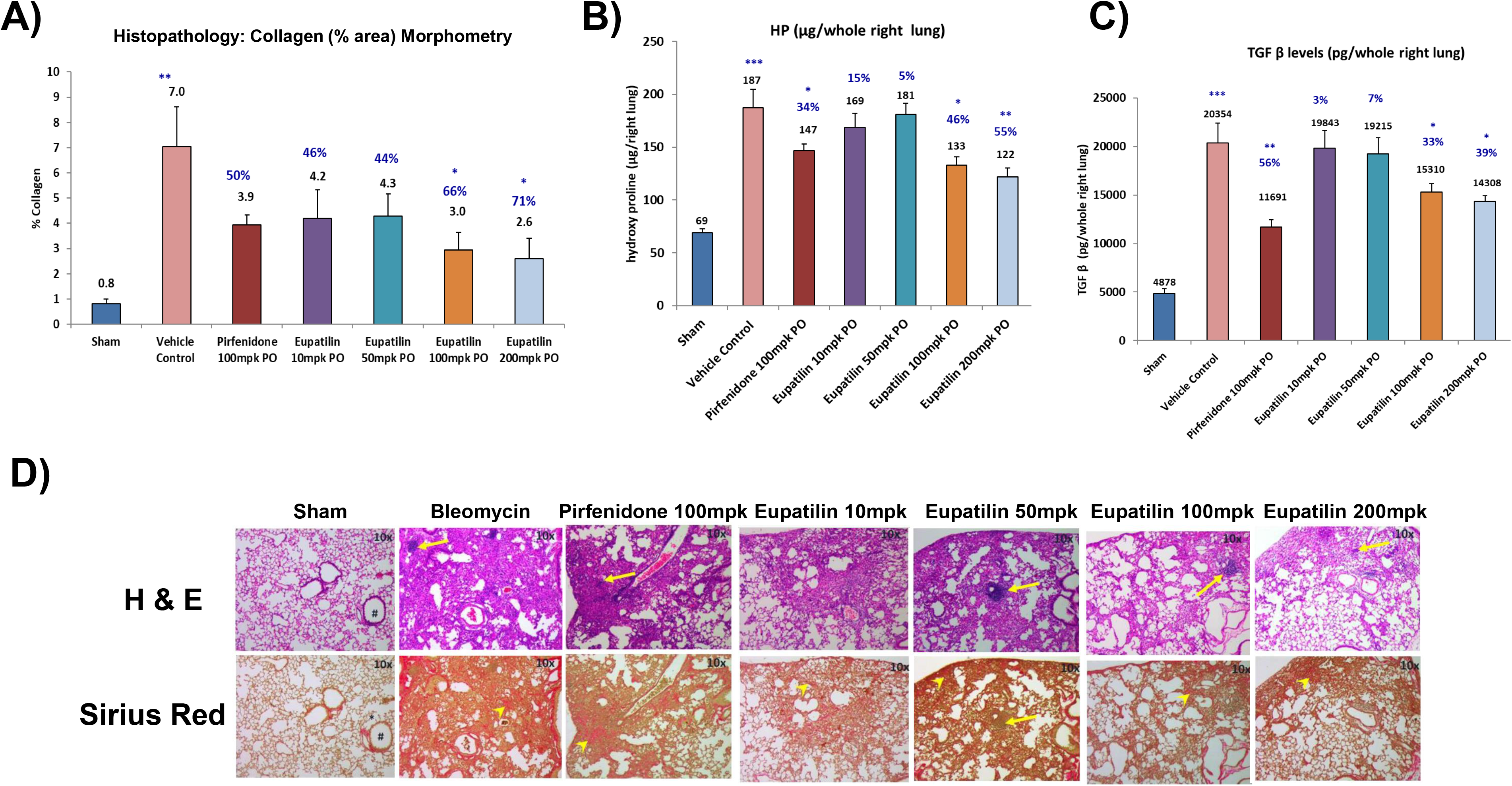
Administration of eupatilin to bleomycin-treated mice significantly improves lung fibrosis. (A) Vehicle (0.5% CMC + 1% Tween 80), pirfenidone (100mpk), or various concentrations of eupatilin (10, 50, 100, or 200mpk) were orally administered into C57BL/6J mice 1 day before administration of bleomycin. Bleomycin was intratracheally injected, and the drugs were administered orally twice per day for 14 days, after which lung tissues and plasma were collected. Histomorphometry was performed. Statistical significance was calculated by Student’s *t*-test (*, p<0.05; **, p<0.01). (B) Levels of hydroxyproline were measured and subjected to Student’s *t*-test as described above. ***, p<0.001. (C) Whole right lung tissue lysates were subjected to a mouse TGFβ ELISA. (D) Paraffin sections of lung tissues were stained with H&E and Sirius Red, and then observed by phase-contrast microscopy (10× magnification).

To determine whether eupatilin has anti-fibrotic efficacy during lung fibrosis, we performed an intervention test using BILF model, as schematized in Figure S10A. Soluble collagen production, collagen histomorphometry and TGFβ production was measured in the lung lysates. We observed significant reductions in collagen levels at eupatilin 100 and 200mpk. As such, pirfenidone 100mpk also had significant anti-fibrotic activity (Figure S10B and S10C), which seemed to be little more efficient than eupatilin in this particular intervention setting. Consistent with the prior therapeutic efficacy BILF test, inhibition of TGFβ production by pirfenidone 100mpk was higher than that of eupatilin 200mpk (Figure S10D). The inhibitory activity by eupatilin 100mpk did not meet a statistical significance. Eupatilin or pirfenidone produced a comparable cell regeneration potential observed by histological staining (Figure S10E). This observation indicates that MOA of eupatilin or pirfenidone may also different *in vivo*, recapitulating the results of the *in vitro* fibrogenesis assays in ONGHEPA1 cells.

### Specific carbons of the CS and chemical residues are critical for anti-fibrogenic effects

The CS of eupatilin, shown by the highlighted carbons in Figure 4A, is linked to a phenyl ring coupled to two methoxy residues. The critical carbon residues for anti-fibrogenic potential are represented in colored fonts: C2 (yellow), C3 (purple), C6 (red) and C7 (green). Next, we selected 34 CDs based on chemical similarities to eupatilin, classified into five chemical groups (A through E), and evaluated their anti-fibrogenic effects (Figure 4E). Among them, three CDs had anti-fibrogenic activity: eupatilin; ONGE200, also known as hispidulin; and ONGA200, also known as jaceosidin. The latter two compounds are the most closely chemically related to eupatilin among the CDs tested. All of these compounds belong to group A, whose members differ in terms of the chemical residues coupled to the phenyl ring. Eupatilin and ONGE200 had marked anti-fibrotic effects (Figure S11A), whereas ONGA300 was cytotoxic at 50 μM (Figure S11B). ONGI300, in which two hydroxy groups are coupled to the phenyl ring, had weak anti-fibrogenesis activity and promoted cell death, suggesting that chemical alteration of the phenyl ring affects anti-fibrotic activity (Figure S11C).

**Figure 4.**
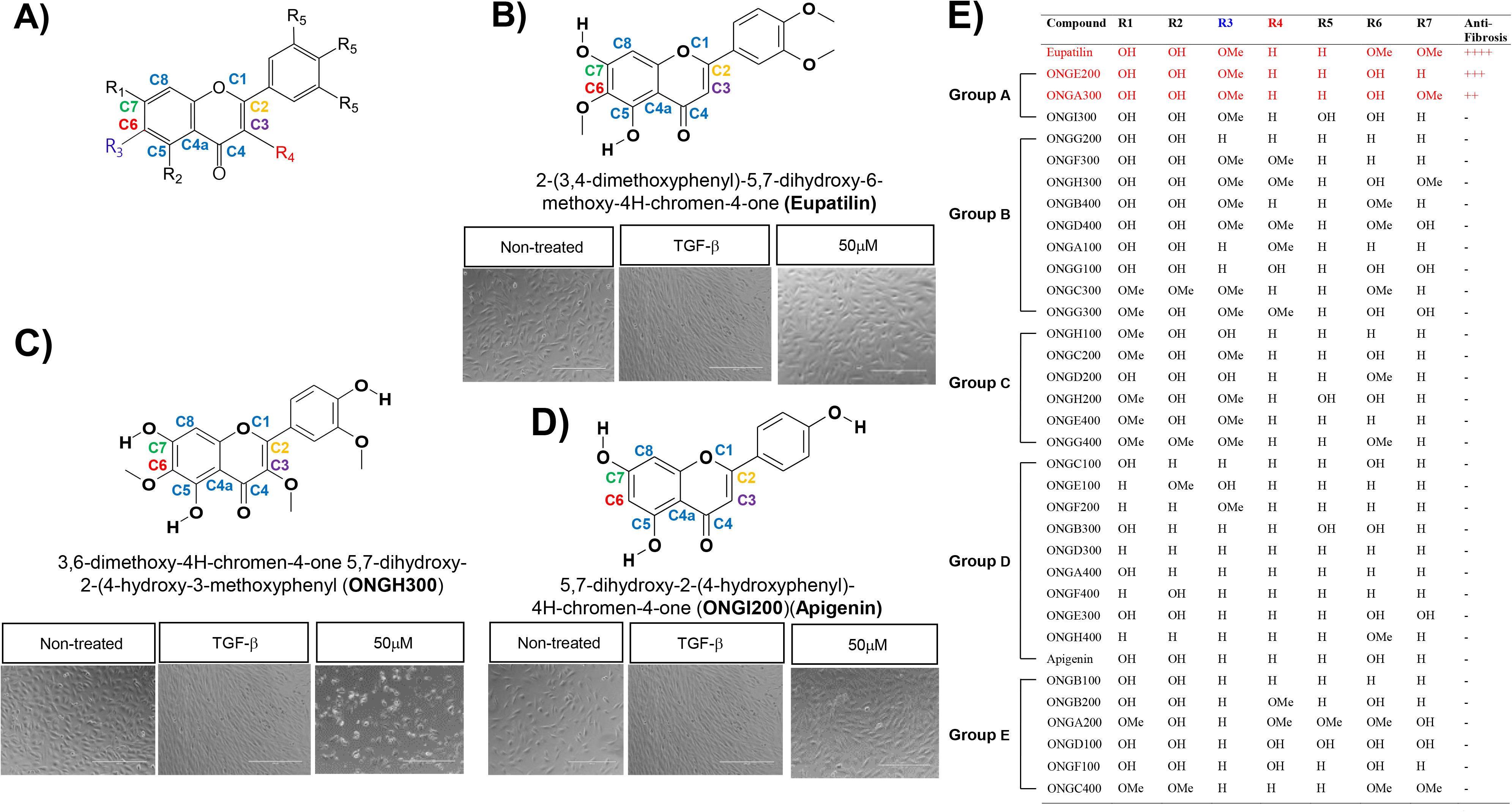
Structure–activity relationship of a selected group of chromone derivatives. (A) Chemical structure of the general chromone scaffold, with its carbons denoted in colored fonts. C2, C3, C6, and C7 are critical for anti-fibrogenesis. (B) ONGHEPA1 cells were simultaneously stimulated with control medium, TGFβ, or TGFβ plus eupatilin (50 μM). Anti-fibrogenic effects were observed by phase-contrast light microscopy. (C) Anti-fibrogenic effects of ONGH300 were observed by phase-contrast light microscopy. (D) Anti-fibrogenic effects of ONGI200 (also called apigenin) were observed by phase-contrast light microscopy. (E) A list of 35 CDs, grouped by chemical moieties coupled to the chromone scaffolds or the phenyl ring. Group A including eupatilin, ONGA300 (jaceosidin), and ONGH300 (hispidulin) exerted potent anti-fibrogenic capacity.

**Figure 5.**
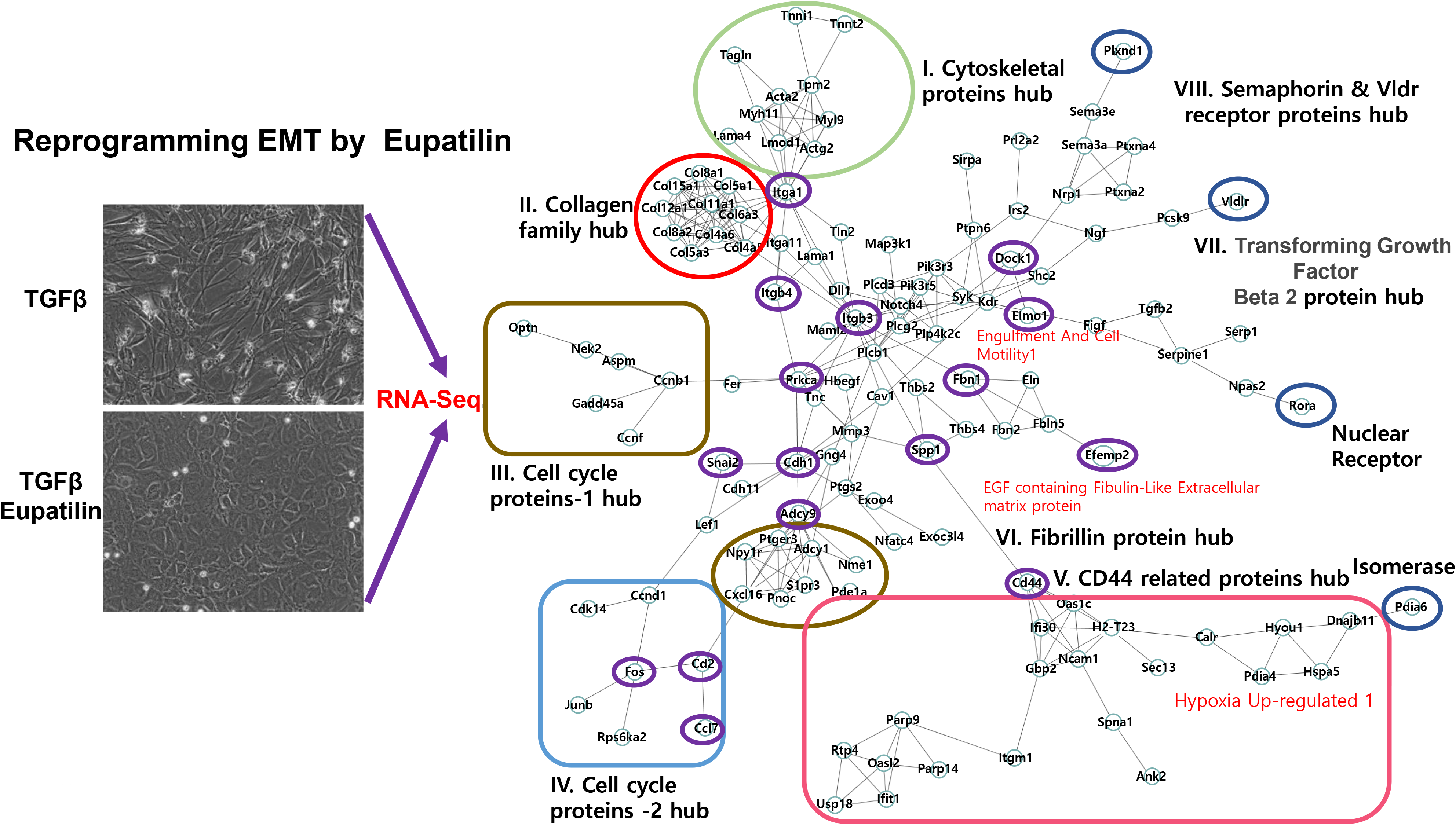
An interactome generated by simultaneous stimulation of ONGHEPA1 cells with TGFβ and eupatilin reveals an EMT-related pathway. Total RNA was isolated from ONGHEPA1 cells treated with TGFβ or TGFβ plus eupatilin. One microgram of RNA from each sample was subjected to RNA-seq, yielding 60 Gb of sequence, and gene interactions were established as described in Experimental procedures. Regulated genes were selected below p<0.05. Eight hubs are represented by colored boxes or ovals, and critical nodes are depicted as purple ovals.

To explore the importance of the eight carbons of the CS in terms of structure–function relationship, two anti-fibrotic CDs (eupatilin and ONGE200) were brought into focus along with the remaining 33 CDs. When a methoxy residue was coupled to C3, as in ONGH300, the anti-fibrotic effect was eliminated and toxicity was apparent (Figure 4C). In comparison with ONGE200, ONGI200 (also known as apigenin) lacks only the methoxy group coupled to C6. ONGI200 had no anti-fibrotic effect, suggesting that C6 is an important constituent of the CS and that methoxylation of this position is necessary to prevent fibrosis (Figure 4D). We also noticed that ONGC200, in which the hydroxy residue at C2 of ONGE200 is replaced with a methoxy group, completely lost the anti-fibrotic effect, suggesting the C7 is also a critical component of the CS (Figure S12A). In addition, methoxylation at C2 of ONGA300, yielding a compound equivalent to ONGH300, abolished anti-fibrotic capacity, suggesting that the presence of a bulky group at C2 may affect anti-fibrogenic potential (Figure S12B). The same held true for ONGD400, a highly cytotoxic stereoisomer of ONGH300 in which C3 is methoxylated (Figure S12C). The chemical residues affecting anti-fibrotic potential are detailed in Figure 4E. Taken together, our data confirm that chemical modifications on the CS are only tolerated at specific carbons, indicating a stringent structure–activity relationship in regard to the anti-fibrogenic effects of these compounds.

### Reprogramming of the EMT by eupatilin may be responsible for anti-fibrosis

We next asked whether eupatilin blocks EMT induced by TGFβ. To this end, we stimulated ONGHEPA1 cells with TGFβ or TGFβ plus eupatilin, and then conducted global gene expression analyses by RNA-seq. Using a bioinformatics approach, we constructed an unbiased interactome that contained several notable hubs, including the collagen gene family (*Col11a1*) and the genes encoding components of the cytoskeleton (*Actin* α *smooth muscle aorta, Troponin 1* and *2*, *Tropomyosin 2, Myosin-heavy chain 1* and *Myosin-light chain 9*, and *Laminin 4*) (Figure S14). Notably, *integrin α1* and *integrin β3* played pivotal roles in connecting these hubs. Expression of *Slug*, a major transcription factor deeply involved in the EMT, was inhibited by eupatilin. Two cell cycle protein hubs, especially *Cdk4* and *c-Fos*, seem to be pronounced, suggesting that the growth arrest effect on DHLFs by eupatilin could be explained. Several genes encoding receptors, including members of the semaphorin receptor family, a cholesterol receptor *(Vldr)*, and *CD44*, formed significant hubs. One notable signaling molecule associated with eupatilin-mediated reprogramming of the EMT might be adenylate cyclase 9 (Adcy9), which is involved in small G-protein activation. To identify the most critical genes involved in reprogramming EMT by eupatilin, we identified a gene set that was maximally induced by TGFβ and maximally repressed by eupatilin. The resultant gene were 103 genes (data not shown). *12*. The vast majority of these genes are involved in promoting EMT. For an instance, follistatin-like 1 *(Fstl1)* plays a pivotal role in pulmonary fibrosis (Dong et al., 2015), and deficiency of *Fstl*1 completely protects mice from bleomycin-induced lung fibrosis, consistent with the downregulation of *Fstl1* by eupatilin. These eupatilin-repressible genes could be new molecular targets for fibrosis.

In conclusion, these observations indicate that eupatilin curbs lung fibrosis by primarily by reprogramming EMT, altering cell shapes and, cell growth arrest and, to a lesser extent than pirfenidone, inhibition of TGFβ production. Therefore, eupatilin could be used for monotherapy or in combinatorial therapy with pirfenidone, with the goal of achieving a synergistic efficacy against IPF.

## Acknowledgments

The authors would like to thank Prof. Ho-Sup Yoon (Nanyang Technological University, Singapore) for insightful comments on the paper. This study was supported by an intramural fund from OsteoNeuroGen.

## Author contributions

B-S Youn and H-S Kim conceived the idea. I-H Kim coordinated the experiments and helped to write the manuscript. J-H Kim was involved in executing major experiments, J-Y Lee contributed to real-time imaging, and Y-M Yoon took part in chemical screening and the EMT assay. MK Maeng conducted phosphorylation analysis following eupatilin stimulation. Mouse care was managed by the IACUC associated with Woojung BSC under IRB 13023.

## Footnotes

The following abbreviated terms were mainly used in this manuscript;

**BILF**: Bleomycin-Induced Lung Fibrosis; **CD**: Chromone Scaffold Derivatives; **CS**: Chromone Scaffold; **DHLFs**: Diseased Human Lung Fibroblasts; **IPF**: Idiopathy Pulmonary Fibrosis; **NASH**:Non-Alcoholic Steato Hepatitis

## Experimental Procedures

### Cell culture and reagents

DHLFs were purchased from Lonza (Basel, Switzerland) and cultured in fibroblast growth medium (FBM, Lonza, Walkersville, MD, USA). Recombinant human TGFβ and PDGF were obtained from Peprotech (Rocky Hill, CT, USA) and used at a final concentration of 5 ng/ml. Chemically synthesized eupatilin was obtained from Syngene International Ltd. (Bangalore, India), dissolved at a stock concentration of 50 mM in DMSO, and stored in aliquots at −20°C. DMSO at 0.1% (v/v) was used as a control. ONGE200 (hispidulin) and ONGA300 (jaceosidin) were purchased from AK Scientific (Union City, CA, USA). The remaining 31 CDs in the test set purchased from Analyticon (Berlin, Germany). For simultaneous treatment, ONGHEPA1 cells (5.5 × 10^4^ cells/well in 1 ml) were seeded in 24-well plates. The anti-fibrotic effect of eupatilin was examined under a microscope (EVOS, Life Technologies, Carsbard, CA, USA).

#### RNA-seq processing, differential gene expression analysis, and interactome analysis

Processed reads were mapped to the *Mus musculus* reference genome (Ensembl 77) using Tophat and Cufflink with default parameters (Trapnell et al., 2012). Differential analysis was performed using Cuffdiff (Trapnell et al., 2012) using default parameters. Further, FPKM values from Cuffdiff were normalized and quantitated using the R Package Tag Count Comparison (TCC) (Sun et al., 2013) to determine statistical significance (e.g., P values) and differential expression (e.g., fold changes). Gene expression values were plotted in various ways (i.e., Scatter, MA, and Volcano plots), using fold-change values, using an R script developed in-house. The protein interaction transfer procedure was performed using the STRING database (Szklarczyk et al., 2011) with the differentially expressed genes. A 60 Gb sequence was generated, and 10,020 transcripts were read and compared. The highest-confidence interaction score (0.9) was applied from the *Mus musculus* species, and information about interacts were obtained based on text mining, experiments, and databases (http://www.string-db.org/).

#### Effects of drugs on bleomycin-induced lung tissue fibrosis

C57BL/6J mice were anesthetized by inhalation of 70% N_2_O and 30% O_2_ gas containing 1.5% isoflurane. Fifty microliters of bleomycin solution in distilled water was directly injected into the lungs, all at once, via the aperture. Immediately after injection, the mice were allowed to recover from the anesthetic, and then housed in normal cages. Bleomycin (0.03U BLM in 50μl saline) was administered once using a visual instillobot. Twelve days after the administration of bleomycin, eupatilin was forcibly nasally administered via a micropipette, once a day (five times a week) for 1 week. Eupatilin was dissolved in DPBS buffer (containing 1% DMSO), and 1 ml/kg was administered based on the most recent body weight. For 2 to 3 days after administration of eupatilin, mice were monitored for toxic symptoms or death, but no abnormal symptoms were observed. Three mice per test group were selected, and their lung tissues were excised. The lung tissues were stained with Masson’s trichrome and observed under a microscope. Results were expressed as mean values and standard deviations. One hour before sacrifice, a final dose of eupatilin or pirfenidone was administered for plasma or lung PK. The bleomycin-treated mice exhibited a rapid decline in weight, but the sham control behaved normally. Eupatilin- or pirfenidone-administered mice exhibited weight gain from day 3 onward. Control and eupatilin-treated mice data were compared using Student’s *t*-test. “Differences between samples were considered statistically significant when p<0.05.

## Supplemental information

### Experimental Procedures

#### Flow cytometry

Cell pellets were suspended in PBS supplemented with 1% FBS and stained with antibodies against CD29, CD44, CD45, CD71, CD90, CD106, and CD117. All antibodies were obtained from BD Biosciences (San Jose, CA, USA). Labeled cells were analyzed on a flow cytometer (FC500, Beckman Coulter, Fullerton, CA, USA).

#### Cell viability assay

Cells were treated with TGFβ and the indicated concentrations of eupatilin (or other CDs) for 24 hr. Cell viability was measured using the CCK-8 kit (Dojindo Molecular Technologies, Tokyo, Japan).

#### Immunostaining

For immunofluorescence staining, ONGHEPA1 cells were grown on 8-well glass chamber slides (LabTek II, Thermo Fisher, Rochester, NY, USA) and fixed with methanol at −80°C for 2 hr. After washing with PBS, cells were blocked and permeabilized by incubation at room temperature for 2 hr in PBS/5% FBS with 0.1% Triton X-100. After incubation with primary antibodies at 4°C overnight, cells were washed, stained with dye-conjugated secondary antibodies, and incubated at room temperature for 2 hr. GATA4 (Santa Cruz Biotechnology, Dallas, TX, USA) and cytokeratin 18 (Santa Cruz Biotechnology) were used as markers of endoderm and endothelial cells, respectively. Nuclei were counterstained with DAPI.

#### Reverse transcriptase PCR and real-time PCR

Cells cultured in 24-well plates were harvested with Trypsin-EDTA solution (Welgene, Seoul, Korea). Total RNA was purified from the pellets using the RNeasy mini Kit (Qiagen, Valencia, CA, USA). RNA was reverse-transcribed using the cDNA Synthesis Kit (PCRBio Systems, London, United Kingdom). Synthesized cDNA was amplified with StepOne Plus (Applied Biosystems, Life Technologies) and 2× qPCRBio Probe Mix Hi-ROX (PCRBio). Comparisons between mRNA levels were performed using the ΔΔsCt method, with *Gapdh* as the internal control.

## Figure legends

**Figure S1.**
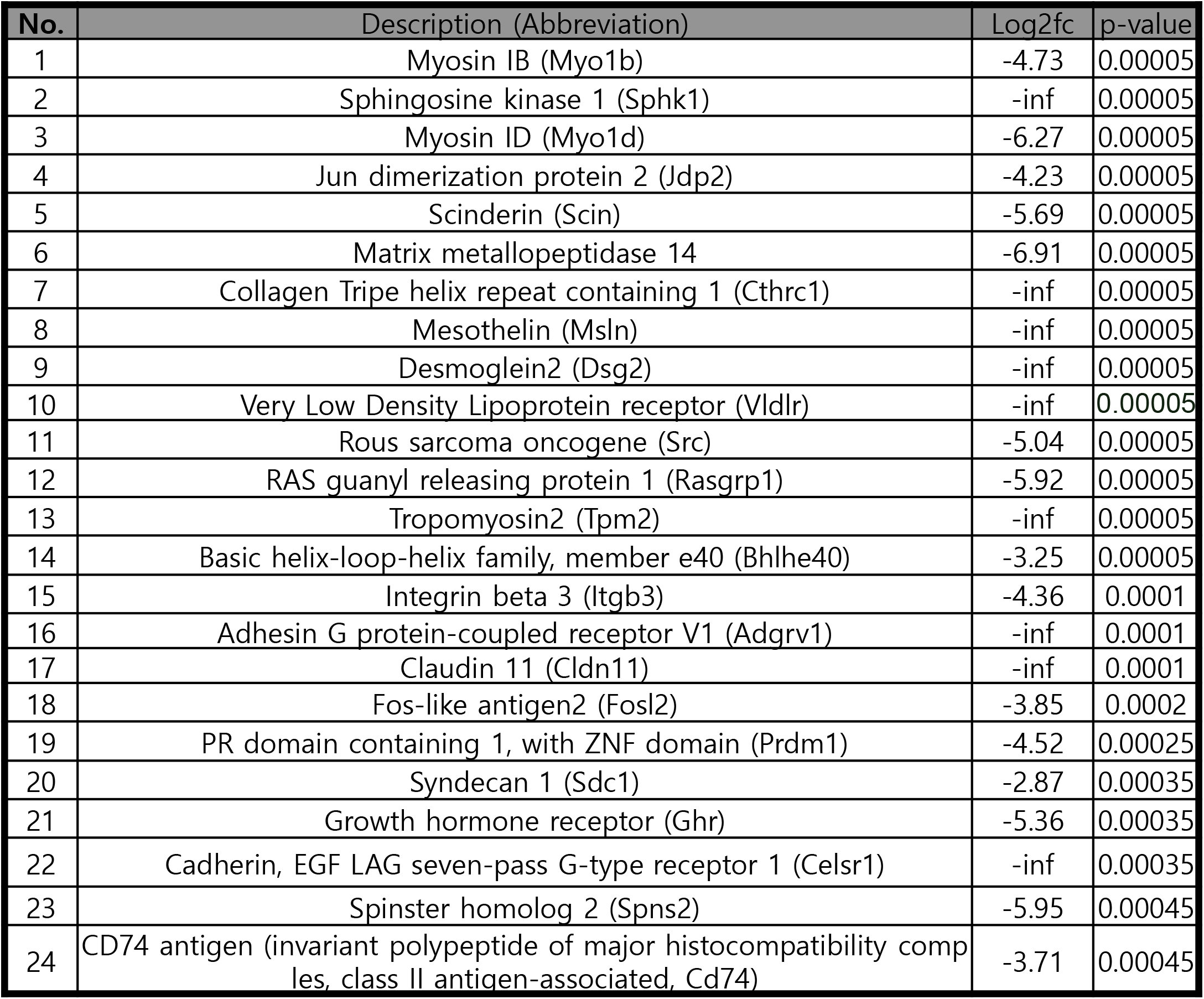
Twenty-four of the top fifty genes most downregulated by eupatilin. Mouse bone marrow cells were prepared and stimulated with M-CSF (20 ng/ml) for 12 hr. Floating cells were collected and re-stimulated with M-CSF for 3 days. Macrophages were stimulated with RANKL (10 ng/ml) in the presence or absence of eupatilin (50 μM). To characterize gene repression by eupatilin, the full transcriptome was analyzed via RNA-seq.

**Figure S2.**
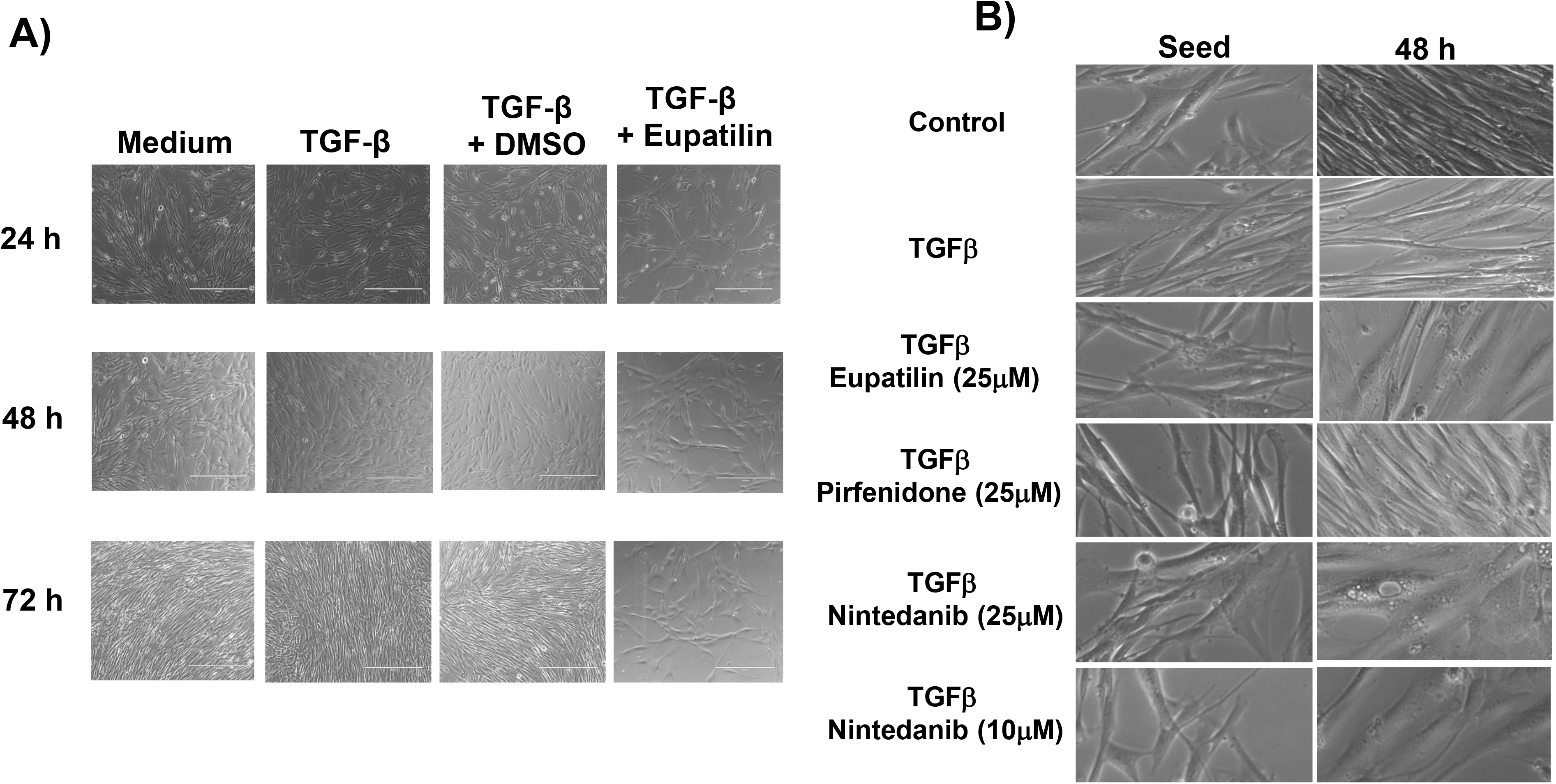
Eupatilin and pirfenidone act differently on DHLFs. (A) DHLFs were seeded in fibroblast growth medium, grown overnight, and then stimulated with control medium, TGFβ, or TGFβ plus eupatilin for 24, 48, or 72 hr. Cell morphology and proliferation were monitored by phase-contrast light microscopy (10×). (B) DHLFs were stimulated with medium, TGFβ, or TGFβ in the presence of eupatilin, pirfenidone, or nintedanib for 48 hr. Morphological change was monitored by phase-contrast light microscopy (100×).

**Figure S3.**
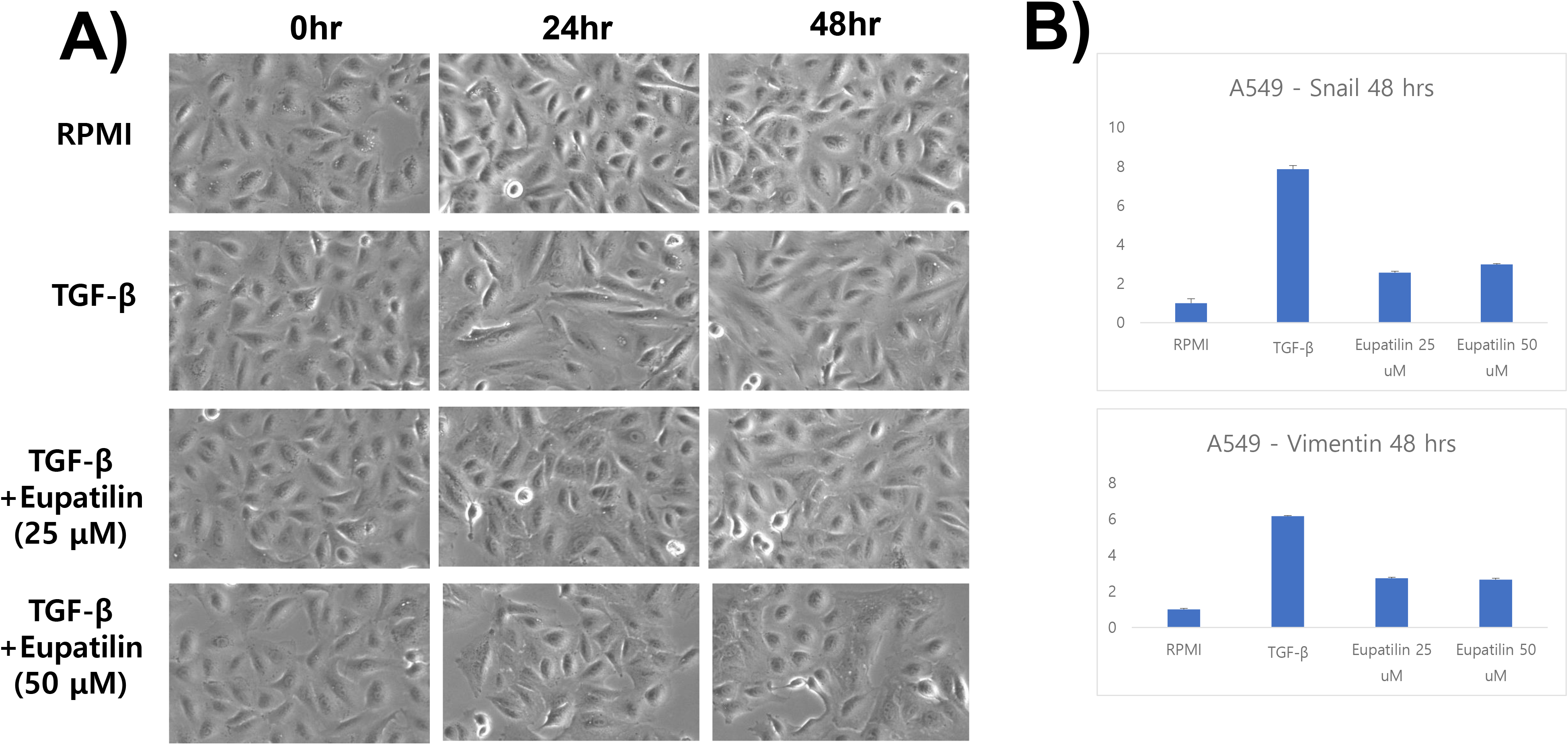
Eupatilin inhibits EMT induction in A549 cells. (A) A549 cells cultured overnight were stimulated with TGFβ or TGFβ plus eupatilin (25 or 50 μM) for the indicated times, and cell shape changes were monitored by phase-contrast microscopy. (B) Total RNA was isolated from A549 cells stimulated with medium, TGFβ, or TGFβ plus eupatilin for 48 hr and subjected to real-time PCR analysis using primers against *snail* or *vimentin*. Standard deviations were calculated from the results of three independent PCRs.

**Figure S4.**
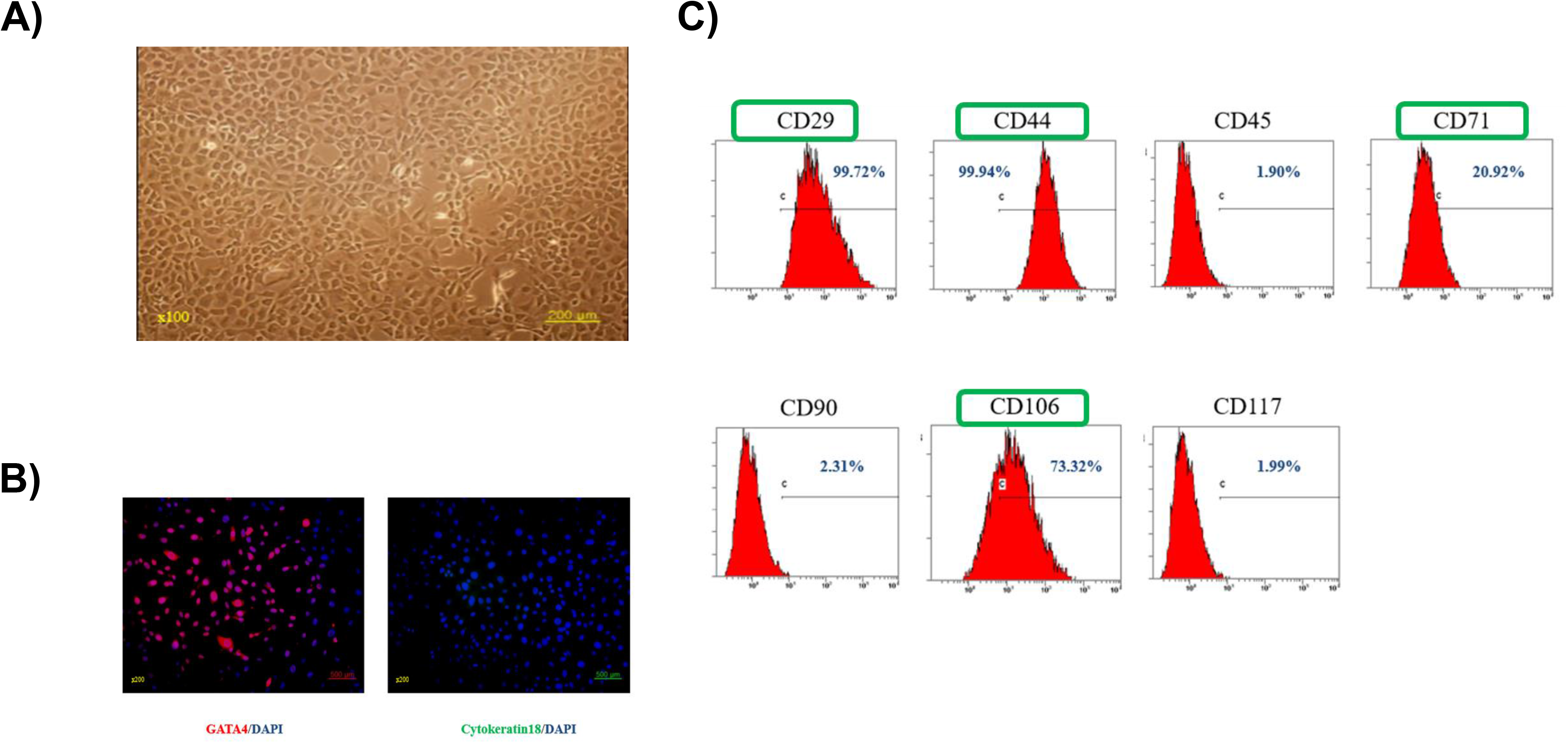
Characterization of ONGHEPA1 cells, a mouse HSC line. (A) Morphology was observed by phase-contrast microscopy at 100× magnification). (B) Immunophenotype of ONGHEPA1. Immunofluorescence staining of GATA4 and Cytokeratin 18. Nuclei were counterstained with DAPI at 200× magnification. (C) Cell surface expression of the CD markers representing HSCs. ONGHEPA1 cells were grown to confluence, washed, and stained with antibodies against CD29, CD44, CD45, CD71, CD90, CD106, and CD117.

**Figure S5.**
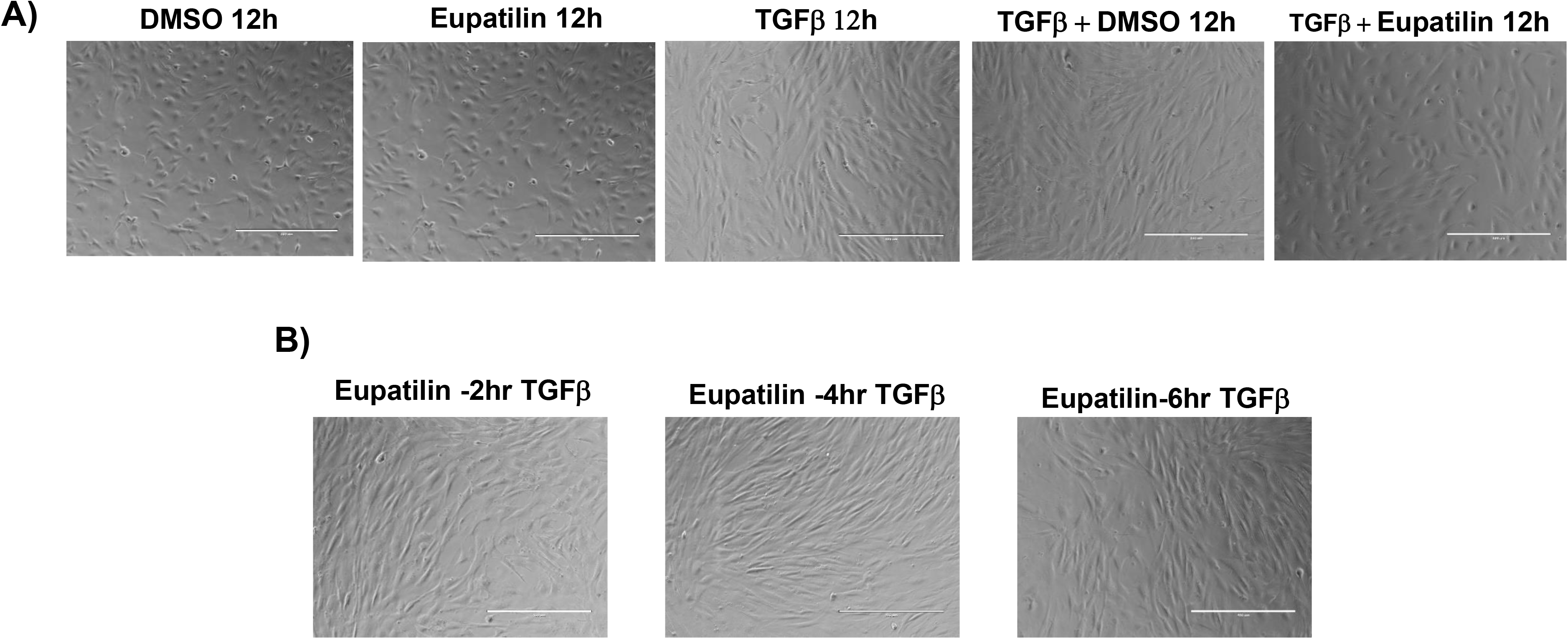
Inhibition of fibrogenesis of ONGHEPA1 cells by eupatilin. (A) ONGHEPA1 cells were differentiated into myofibroblasts in the presence of DMSO, eupatilin, TGFβ, or TGFβ plus eupatilin for 12 hr. Anti-fibrogenic capacities were assessed by phase-contrast light microscopy. (B) ONGHEPA1 cells were pre-stimulated with eupatilin for 2, 4, or 6 hr, washed, and then stimulated with TGFβ for 24 hr. Anti-fibrogenic capacities were assessed by phase-contrast light microscopy.

**Figure S6.**
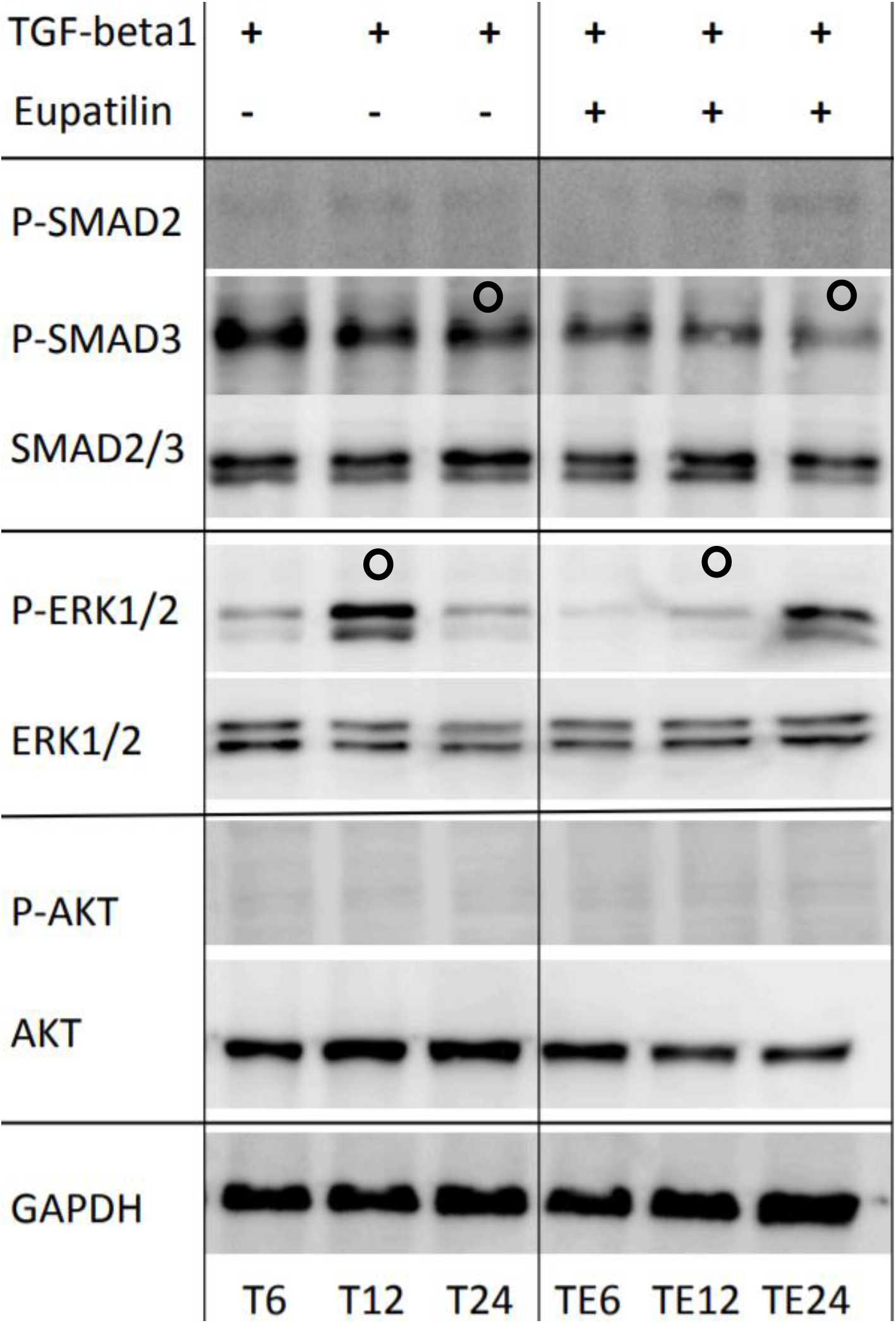
Inhibition of the phosphorylation of Erk and Smad3 by eupatilin. ONGHEPA1 cells were stimulated with TGFβ or TGFβ plus eupatilin for 6, 12, or 24 hr. The treated cells were washed, and cell lysates were prepared for western blot analysis. The membranes were incubated with SMAD2/3 mAb Sampler kit, ERK 1/2 mAb, phospho-ERK 1/2 mAb, AKT mAb, phospho-AKT mAb, or GAPDH mAb. Immunoreactive proteins were visualized by chemiluminescence and recorded with a digital recorder (LAS 4000 mini, Fuji Film, Tokyo, Japan). Decreased phosphorylation of Erk or SMAD3 is denoted by empty circles. T6: ONGHEPA + TGFβ, 6 hr; T12: ONGHEPA + TGFβ, 12 hr; T24: ONGHEPA + TGFβ, 24 hr; TE6: ONGHEPA + TGFβ + Eupatilin, 6 hr; TE12: ONGHEPA + TGFβ + Eupatilin, 12 hr; TE24: ONGHEPA + TGFβ + Eupatilin, 24 hr.

**Figure S7.**
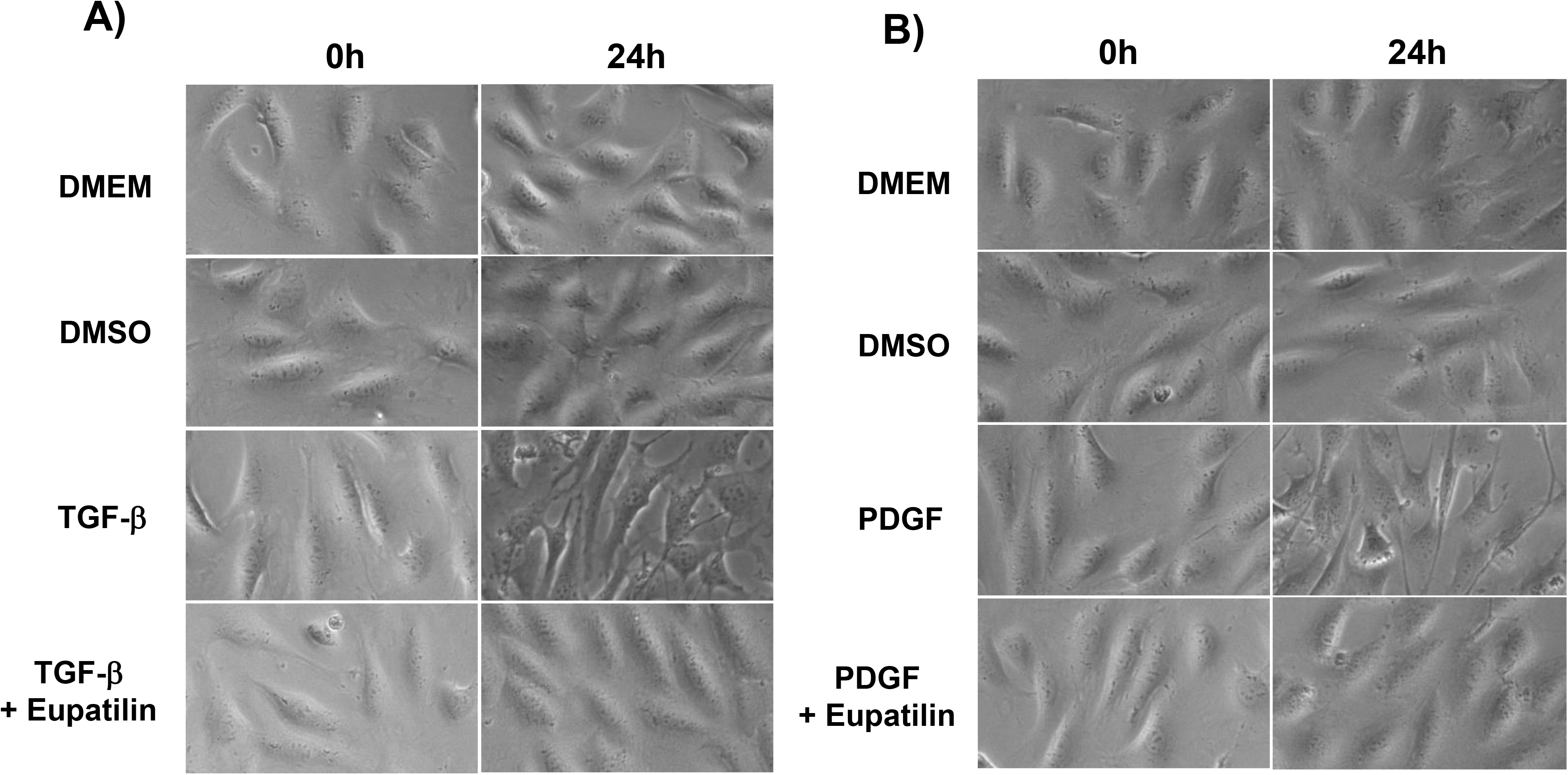
Eupatilin inhibits fibrogenesis in ONGHEPA1 cells stimulated with TGFβ or PDGF. (A) ONGHEPA1 cells were stimulated with medium, DMSO, TGFβ, or TGFβ plus eupatilin for 24 hr. Morphological changes were monitored by phase-contrast light microscopy (100×). (B) ONGHEPA1 cells were stimulated with medium, DMSO, PDGF, or PDGF plus eupatilin for 24 hr. Morphological changes were monitored by phase-contrast light microscopy (100×).

**Figure S8.**
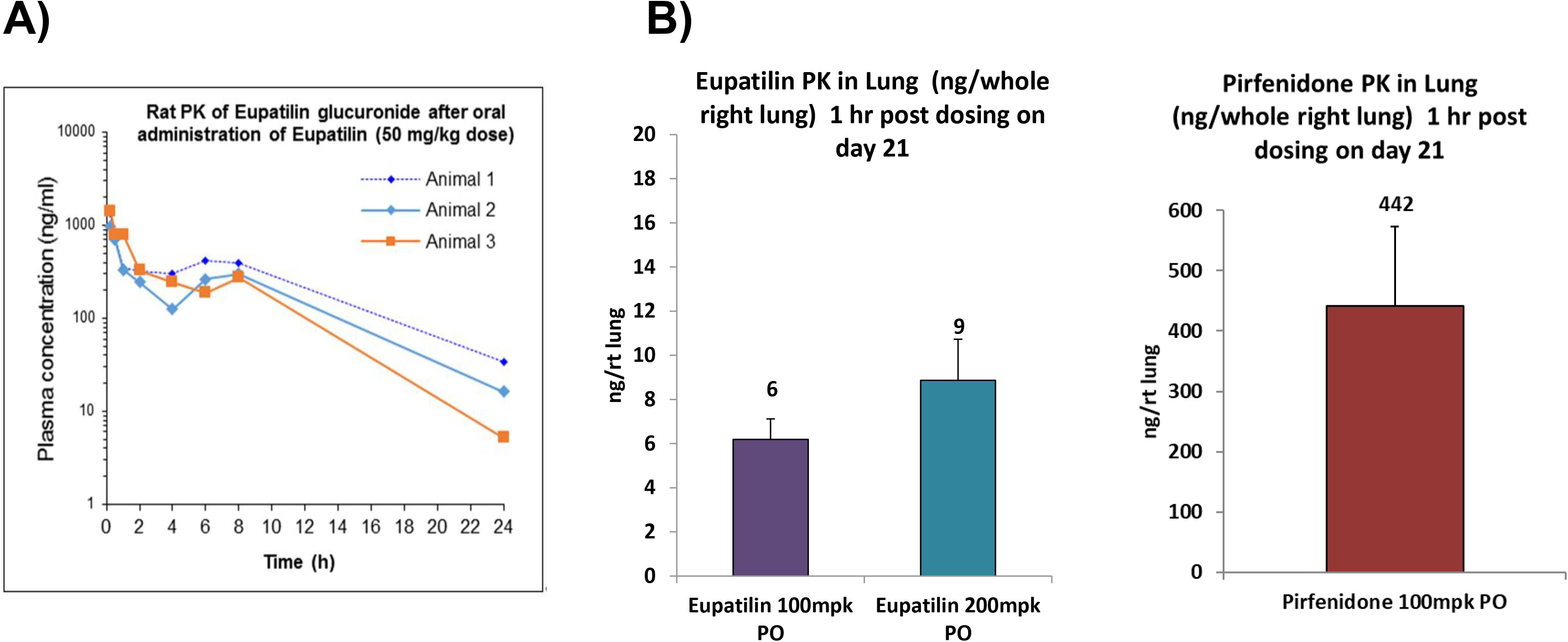
Comparison of PK between eupatilin and pirfenidone. (A) Three rats were orally administered 50mpk eupatilin, and plasma was collected at various time points. Concentrations of parental eupatilin or glucuronide-eupatilin were determined by HPLC. (B) Rats were orally administered 100mpk eupatilin, 200mpk eupatilin, or 100mpk pirfenidone. Protein lysates from whole right lungs were prepared, and levels of parental eupatilin or pirfenidone were determined by HPLC.

**Figure S9.**
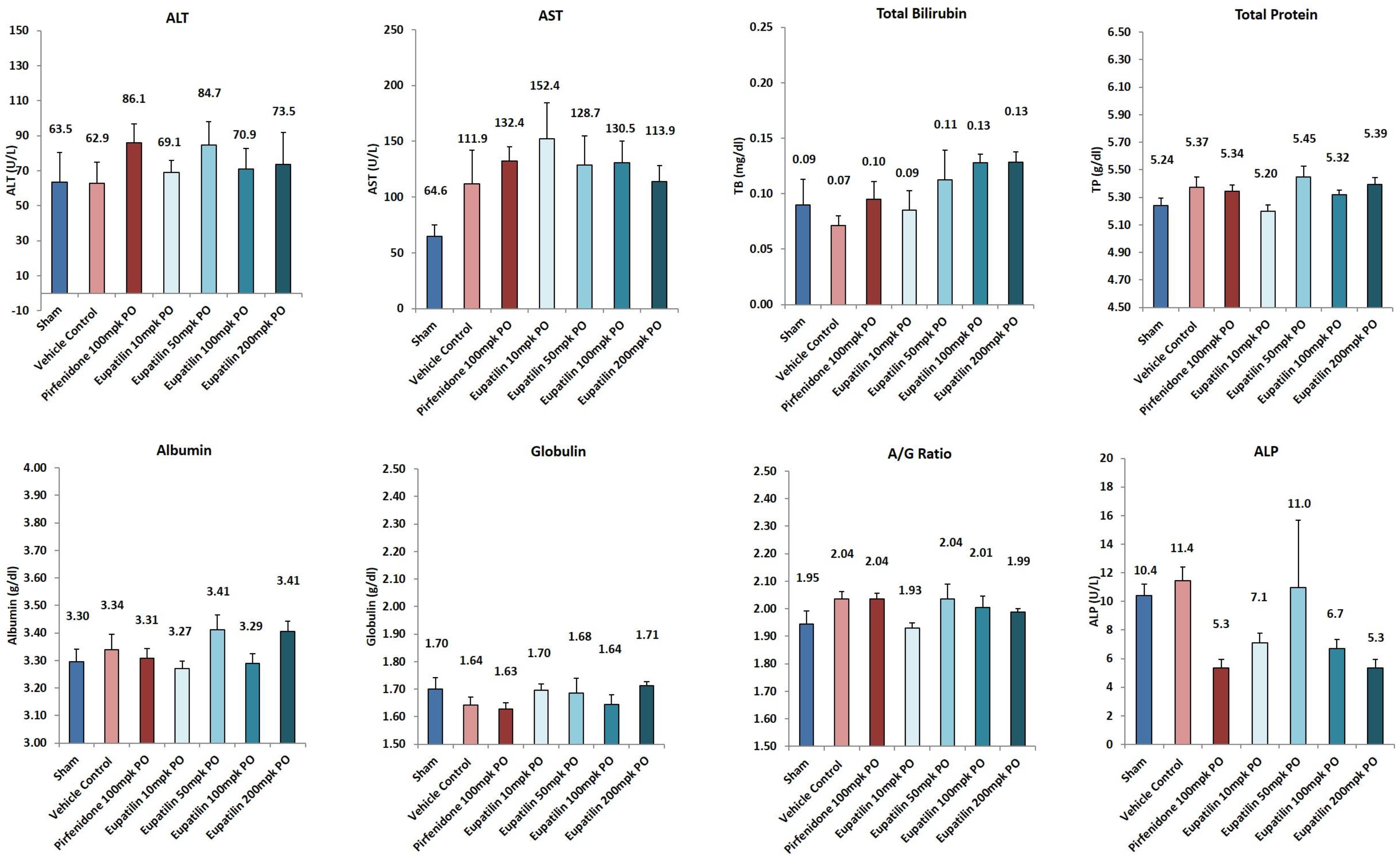
Eupatilin exhibited no hepatotoxicity. Mice were treated with vehicle alone, pirfenidone (100mpk), or eupatilin (10, 50, 100, or 200mpk) in conjunction with intratracheal administration of bleomycin. Plasma samples were collected and subjected to liver function tests (ALT, AST, ALP, total bilirubin, albumin, and globulin).

**Figure S10.**
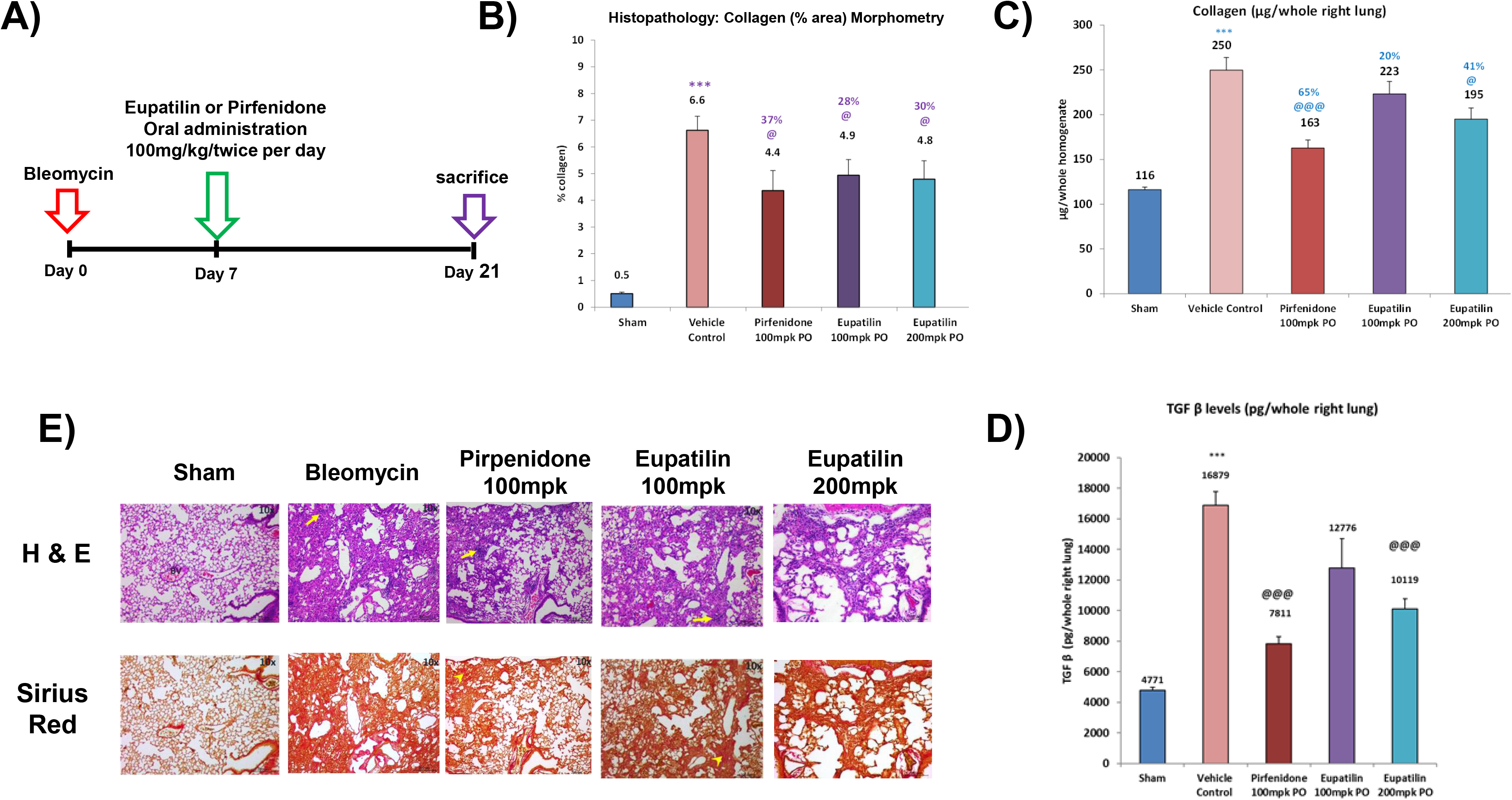
Amelioration of lung fibrosis by eupatilin. (A) Bleomycin was intratracheally administered into mice at day G, and lung fibrosis was generated for 7 days. Eupatilin (100 or 200mpk) or pirfenidone (100mpk) was orally administered for 14 days. Lung lysates and tissue paraffin sections were prepared. (B) Histomorphometry was examined, and statistical significance was calculated by Student’s *t*-test; *, p<0.05. (C) Soluble collagen from whole right lung was measured. *, p<0.05 (Student’s *t*-test). (D) TGFβ in whole right lung was measured by ELISA. ***, p<0.001 (Student’s *t*-test). (E) Paraffin sections of lung tissues were stained with H&E or Sirius Red and examined using phase-contrast microscopy at 4× or 20× magnification.

**Figure S11.**
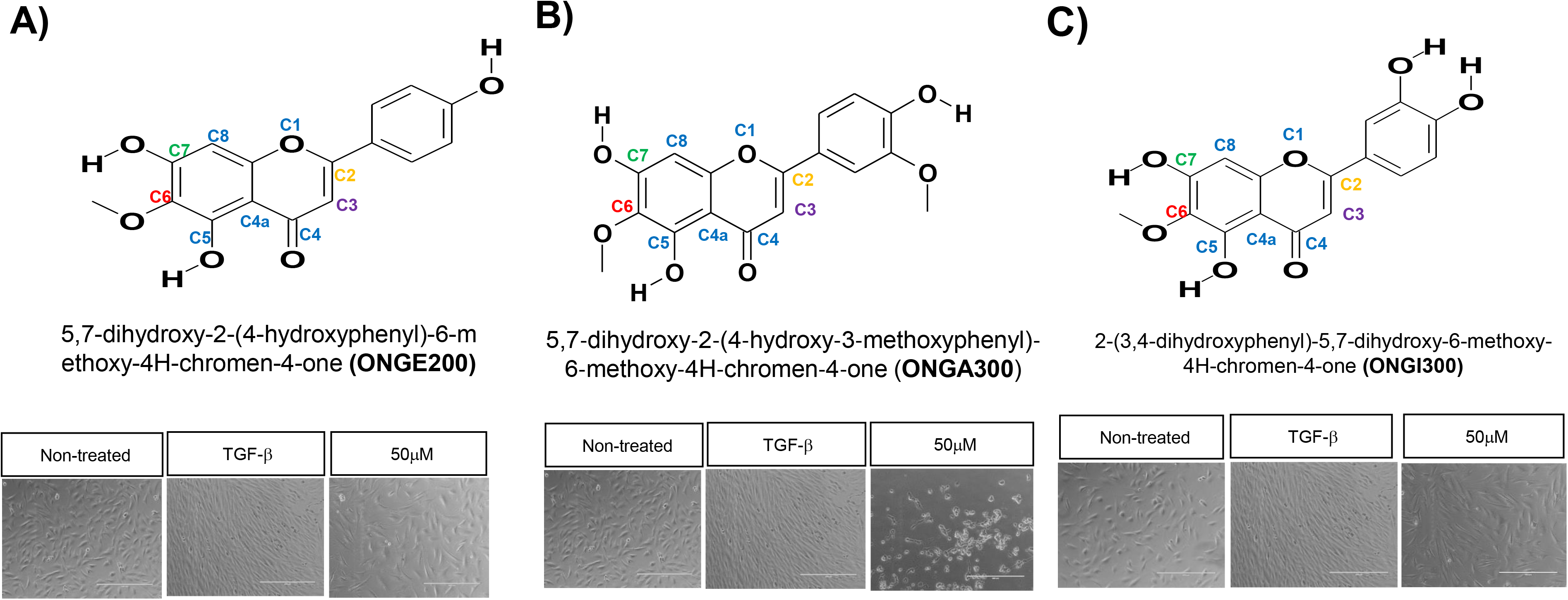
Structure-activity relationship of a selected group of chromone derivatives with minor chemical moieties coupled to the phenyl ring of eupatilin. The structure–activity relationships of chromone compounds with similar chemical structures are summarized in the text (Fig. 4). ONGHEPA1 cells were treated with TGFβ for 24 hr in the presence or absence of the indicated compounds. Anti-fibrotic activities were monitored by light microscopy. Three chromones (ONGE200, ONGA300, and ONGI200) exhibited no or extremely low anti-fibrotic activities.

**Figure S12.**
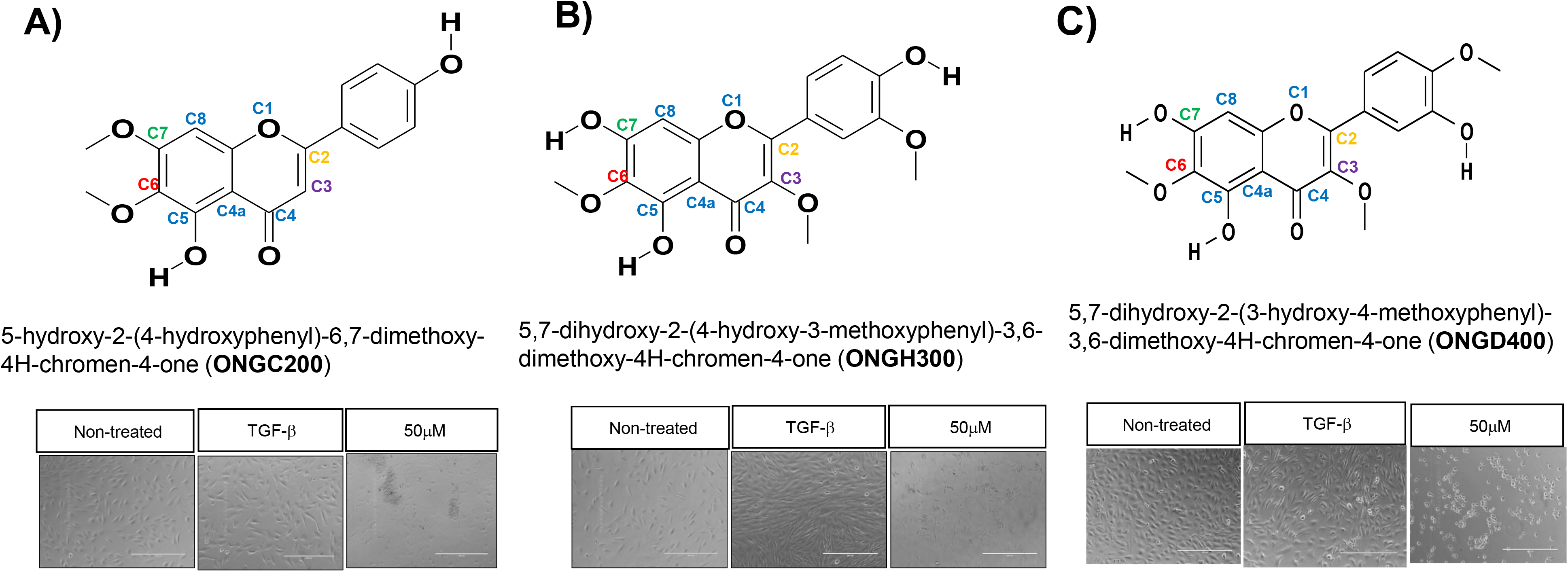
Structure–activity relationship of a selected group of chromone derivatives with different chemical moieties coupled to the chromone scaffold of eupatilin. ONGHEPA1 cells were treated with TGFβ for 24 hr in the presence or absence of the indicated compounds. Anti-fibrotic activities were monitored by light microscopy. Three chromones (ONGC200, ONGH300, and ONGD400) exhibited no anti-fibrotic activities.

